# A Bayesian approach to inferring dispersal kernels with incomplete mark-recapture data

**DOI:** 10.1101/813790

**Authors:** Akira Terui

## Abstract

Dispersal is a fundamental ecological process that links populations, communities and food webs in space. However, dispersal is tremendously difficult to study in the wild because we must track individuals dispersing in a landscape. One conventional method to measure animal dispersal is a mark-recapture technique. Despite its usefulness, this approach has been recurrently criticized because it is virtually impossible to survey all possible ranges of dispersal in nature. Here, I propose a novel Bayesian model to better estimate dispersal parameters from incomplete mark-recapture data. The dispersal-observation coupled model, DOCM, can extract information from both recaptured and unrecaptured individuals, providing less biased estimates of dispersal parameters. Simulations demonstrated the usefulness of DOCM under various sampling designs. I also suggest extensions of the DOCM to accommodate more realistic scenarios. Application of the DOCM may, therefore, provide valuable insights into how individuals disperse in the wild.

## Introduction

Ecological entities rarely exist independently. Dispersal – any movement of organisms across space – links populations (Hanski 1999; Hanski & Ovaskainen 2000; Terui *et al.* 2018a; Terui *et al.* 2014b), communities (Leibold *et al.* 2004; Terui & Miyazaki 2016) and food webs (Nakano *et al.* 1999; Nakano & Murakami 2001; Spiller *et al.* 2010; Terui *et al.* 2018b) that are otherwise isolated from one another. Ecologists have been intrigued by the spatial process due to its implications for critical applied issues, such as the metapopulation persistence of endangered species in fragmented landscapes (Hanski 1999). More recently, accumulating evidence suggests that dispersal is highly plastic and context-dependent with significant consequences for landscape-level dynamics (Bonte & de la Pena 2009; Bonte *et al.* 2012; Cote *et al.* 2011; Cote *et al.* 2013; Fronhofer *et al.* 2018; Fronhofer *et al.* 2017; Little *et al.* 2019; Terui *et al.* 2017). For example, Fronhofer *et al.* (2018) have shown that context-dependent dispersal in experimental landscapes has stabilizing effects on local food webs coupled via dispersal. Therefore, there is an increasing awareness that an in-depth understanding of dispersal is critical to biodiversity forecasting during rapid environmental changes. Nevertheless, dispersal is inherently difficult to study in the wild (Nathan 2001). There have been many attempts to track dispersal in natural systems (Clobert *et al.* 2012; Comte & Olden 2018; Nathan *et al.* 2008), but linking dispersal processes with specific ecological factors has been challenged by the incomplete nature of field observations. As such, a detailed analysis of dispersal is, to some extent, biased towards small-scale controlled experiments, limiting our ability to infer spatial dynamics at large spatial scales.

One conventional method to measure animal dispersal in the wild is a mark-recapture technique. Although mark-recapture studies can provide valuable insights into how individuals move across space, there are some serious problems when applying this method in nature. First, it is virtually impossible to survey all possible range of dispersal in a landscape (Fujiwara *et al.* 2006; Gowan & Fausch 1996; Schwalb *et al.* 2010; Terui *et al.* 2014a). Consequently, a substantial portion of individuals can leave behind the study area, causing serious underestimation of dispersal parameters. Second, even when marked individuals remained in the study area, imperfect detection of marked individuals may pose a challenge to infer dispersal processes (Pepino *et al.* 2012; Rodriguez 2002). To date, several statistical models have been proposed to overcome these difficulties (Fujiwara *et al.* 2006; Pepino *et al.* 2012; Rodriguez 2002). For example, Rodriguez (2002) developed a general class of dispersal models that describe how marked individuals are recaptured through dispersal and sampling processes. However, these models are implicit about unrecaptured individuals and/or have limited extendibility to more complex models that capture plastic and context-dependent dispersal. Hence, there is a need to develop a new class of statistical models that have a greater extension capacity.

Bayesian inference provides a flexible statistical framework that may open the opportunity to overcome challenges in utilizing mark-recapture data (Kéry & Schaub 2012; Terui *et al.* 2017). Here, I introduce a novel Bayesian model that integrates dispersal and observation processes into a single coupled model. The dispersal-observation coupled model, DOCM, can extract information from both recaptured and unrecaptured individuals. Consequently, the model can provide less biased estimates of ecological parameters. In this study, I demonstrate that the usefulness of DOCM using simulated test datasets produced under various sampling designs and discuss its extension capacity to more realistic models.

## Model

I consider a situation in which a virtual ecologist conducts a mark-recapture study in a one-dimensional space (e.g., a stream). They choose a section with length *L* for the mark-recapture study (i.e., the observation section) and divide it into subsections with length *l*. The number of subsections is thus *L l ^−1^*. In each subsection, virtual ecologists perform an initial capture survey and assign a subsection ID to each individual to locate them. After marking individuals uniquely, captured individuals are released into the center of the subsection where they were caught. Then, released individuals disperse freely for a certain period and a recapture survey occurs in the observation section. Since the observation section is a finite domain, individuals can leave this area. Also, only survived individuals may be recaptured with some probability even when marked individuals stay in the observation section. Thus, to be recaptured, individuals must (1) stay in the observation section, (2) survive until being recaptured, and (3) be detected if they survive and remain in the observation section. To represent this data-producing process, I propose the following modeling framework that integrates dispersal and observation processes (Figure 1).

**Figure 1.**
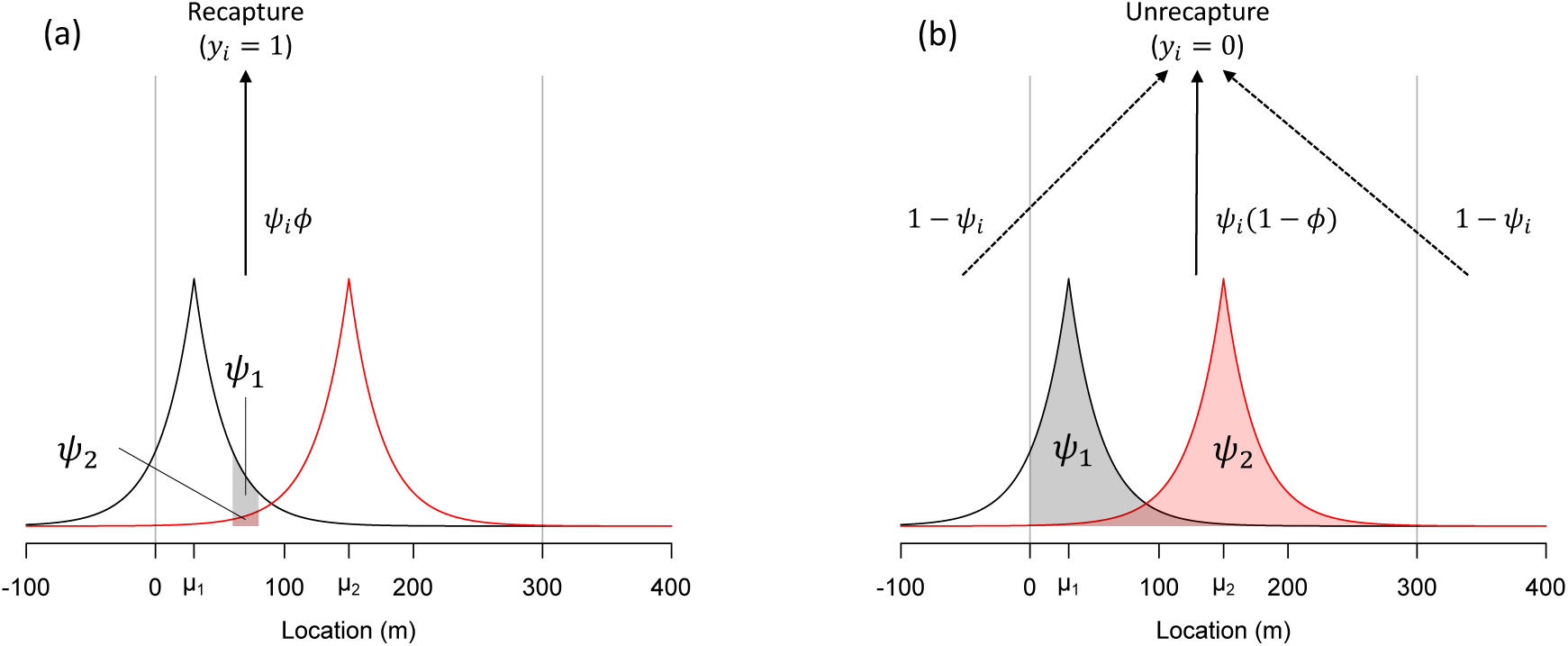
Graphical representation of the dispersal-observation coupled model (DOCM). The black and red lines are the examples of Laplace dispersal kernels for individual 1 and 2 released at different locations (average dispersal distance *δ* = 25 m for both kernels). Vertical gray lines indicate the observation section (0 – 300 m for this example). Shaded areas denote *ψ*_*i*_ that represents the probability that an individual moves from the release location *μ*_*i*_ to the recapture subsection for recaptured individuals (a) or the probability that an individual stays in the observation section for unrecaptured individuals (b). Individuals released at different locations (*μ*_1_ and *μ*_2_) have different values of *ψ*_*i*_. After the dispersal process, individuals are subject to incomplete recapture surveys, by which individuals may be detected with the recapture probability *ϕ* if they remained in the observation section. Note that the recapture probability *ϕ* is a composite of survival and detection probabilities.

### Dispersal model

I first model the dispersal process. Let *μ*_*i*_ and *x*_*i*_be locations at initial capture and recapture sessions, respectively, for individual *i*. The variables *μ*_*i*_ and *x*_*i*_may be expressed as the distance from the center of the capture/recaptured subsection to either end of the observation section (e.g., the downstream end of the study section in streams). I assume the location variable at recapture *x*_*i*_to follow a Laplace distribution, a dispersal kernel commonly used in the dispersal literature (Nathan *et al.* 2012; Rodriguez 2002):

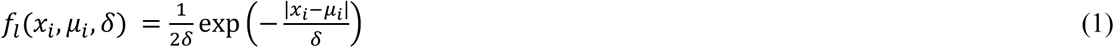

The parameter *δ* is the average dispersal distance. Equation 1 illustrates that the recapture location *x*_*i*_is conditional on the release location *μ*_*i*_ and the dispersal parameter *δ* (Figure 1).

### Observation model

After dispersal, marked individuals are subject to an imperfect observation process. Let *y*_*i*_ be the variable representing a recapture history for individual *i* (*y*_*i*_ = 1 if recaptured; otherwise 0). The response variable *y*_*i*_ can be modeled as a realization of a Bernoulli distribution:

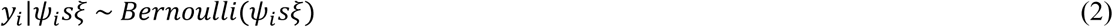

where *ψ*_*i*_ is the probability that individual *i* moves to the subsection of recapture (recaptured individuals) or stays in the observation section (unrecaptured individuals), *s* is the survival probability between the time points of release and recapture, and *ξ* is the detection probability during a recapture survey. The parameters *s* and *ξ* can be isolated if an independent dataset to estimate detection probability, e.g., multiple-pass removal data, is available (Dorazio *et al.* 2005). Otherwise, the two parameters need to be condensed into recapture probability *ϕ* (=*sξ*) such that:

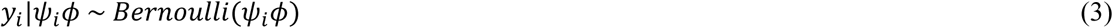

Here, I couple the observation and dispersal models by describing *ψ*_*i*_ as a function of the dispersal parameter *δ* and release location *μ*_*i*_. Specifically, *ψ*_*i*_ is denoted as:

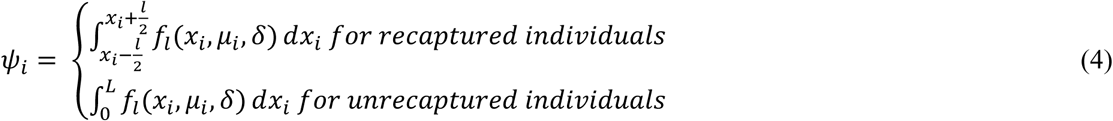

Recaptured individuals are known to be present at the subsection of recapture, so the range of integration is given as 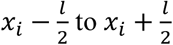 in equation 4 (i.e., from one end to another end of the subsection). This expression gives the probability of movement from the release location *μ*_*i*_ to the subsection of recapture given the dispersal parameter *δ*. For unrecaptured individuals, equation 4 accounts for two important facts: (1) a greater value of the dispersal parameter decreases the probability of remaining in the observation section (*ψ*_*i*_) and (2) the release location *μ*_*i*_ influences *ψ*_*i*_ (i.e., individuals released near the edge of the observation section are more likely to emigrate; Figure 1b). Key parameters in the DOCM were summarized in Table 1.

**Table 1.** Key parameters used in the dispersal-observation coupled model (DOCM)

| Parameter | Interpretation |
| --- | --- |
| $\delta$ | Average dispersal distance |
| $\mu$ | Release location |
| $\psi$ | Probability of moving from the release location $\mu$ to the recapture subsection (recaptured individuals); Probability of remaining in the observation section (unrecaptured individuals) |
| $\phi$ | Recapture probability |
| $s$ | True survival probability |
| $\xi$ | Detection probability |

### Evaluation of model performance

To evaluate the performance of the DOCM, I generated test datasets under different sampling designs. Specifically, I focused on the following design factors that are related to sampling efforts in the field: (1) the number of individuals marked and released *N* (100, 500, and 1000 individuals), (2) the length of the observation section *L* (500 and 1000 m) and (3) the length of an individual subsection or resolution *l* (20 and 50 m) (Figure 2). In addition, I considered variation in the recapture probability *ϕ* (0.25, 0.50, and 0.75). I considered all possible combinations of *N, L, l*, and *ϕ* (36 combinations) when generating test datasets.

**Figure 2.**
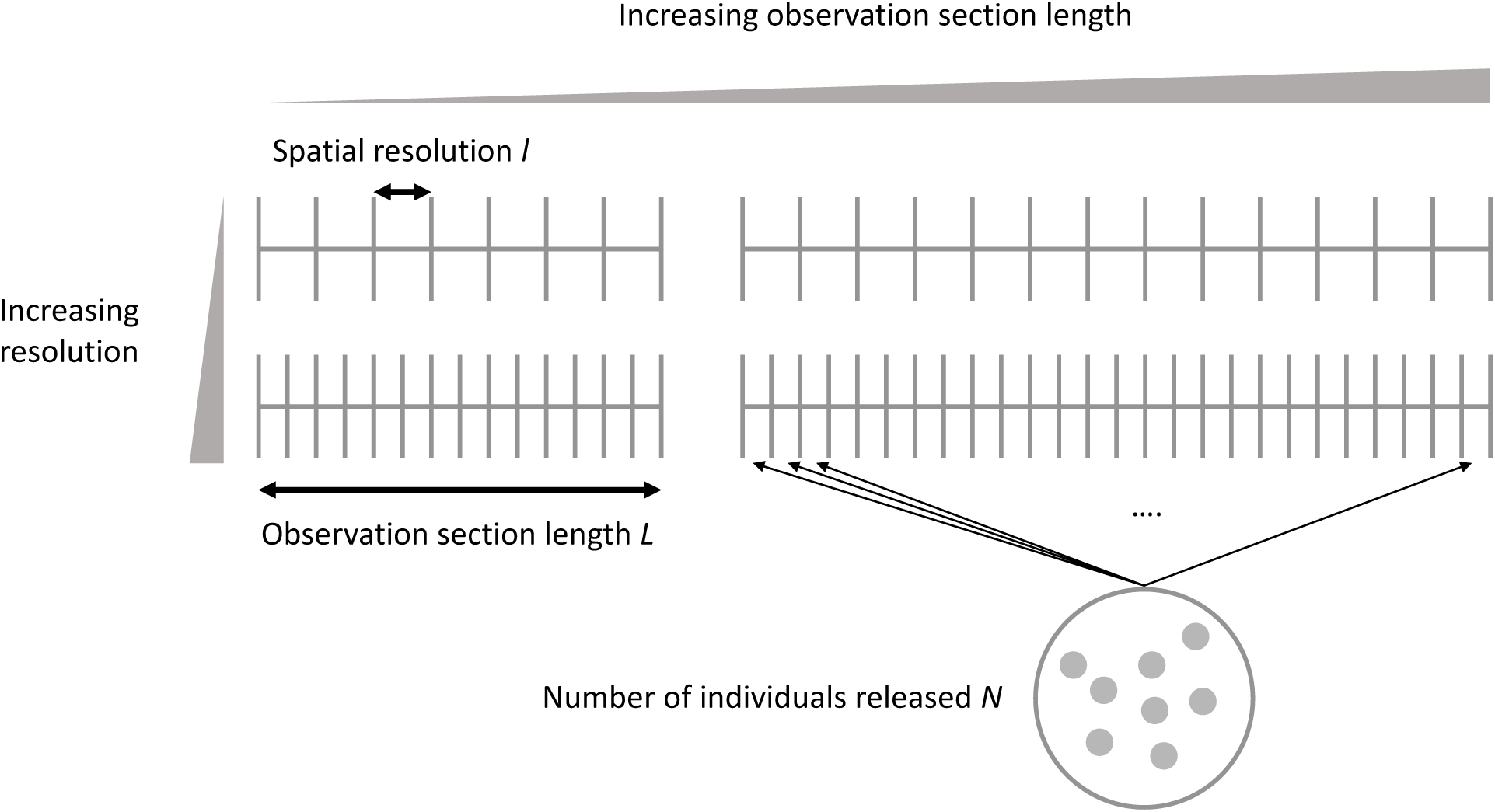
Schematic diagram of sampling designs. Three design factors were considered: (1) the number of individuals released *N*, (2) observation section length *L*, and (3) spatial resolution *l*.

Under each sampling design, I produced 100 test datasets with different values of *δ*, which was drawn from a uniform distribution (range: 10 – 300 m). Each independent dataset was generated as follows. First, *N* individuals were assigned randomly to 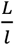 subsections (Figure 3a). These “marked” individuals were released at the center of the captured subsection, which was recorded as release location *μ*_*i*_. Second, released individuals relocate themselves along a one-dimensional space according to a known dispersal kernel as *x*_*i,true*_| *μ*_*i*_, *δ ~ Laplace*(*μ*_*i*_, *δ*) (Figure 3b). Individuals were considered to remain in the observation section if true recapture location *x*_*i,true*_ was within a range of 0 – *L* m. Then, remained individuals were recaptured with recapture probability *ϕ* (Figure 3c). When recaptured, the true recapture location *x*_*i,true*_was rounded to a location value at the center of the recapture subsection *x*_*i*_to mimic real field data (Figure 3c). For unrecaptured individuals, *x*_*i*_was recorded as “NA”.

**Figure 3.**
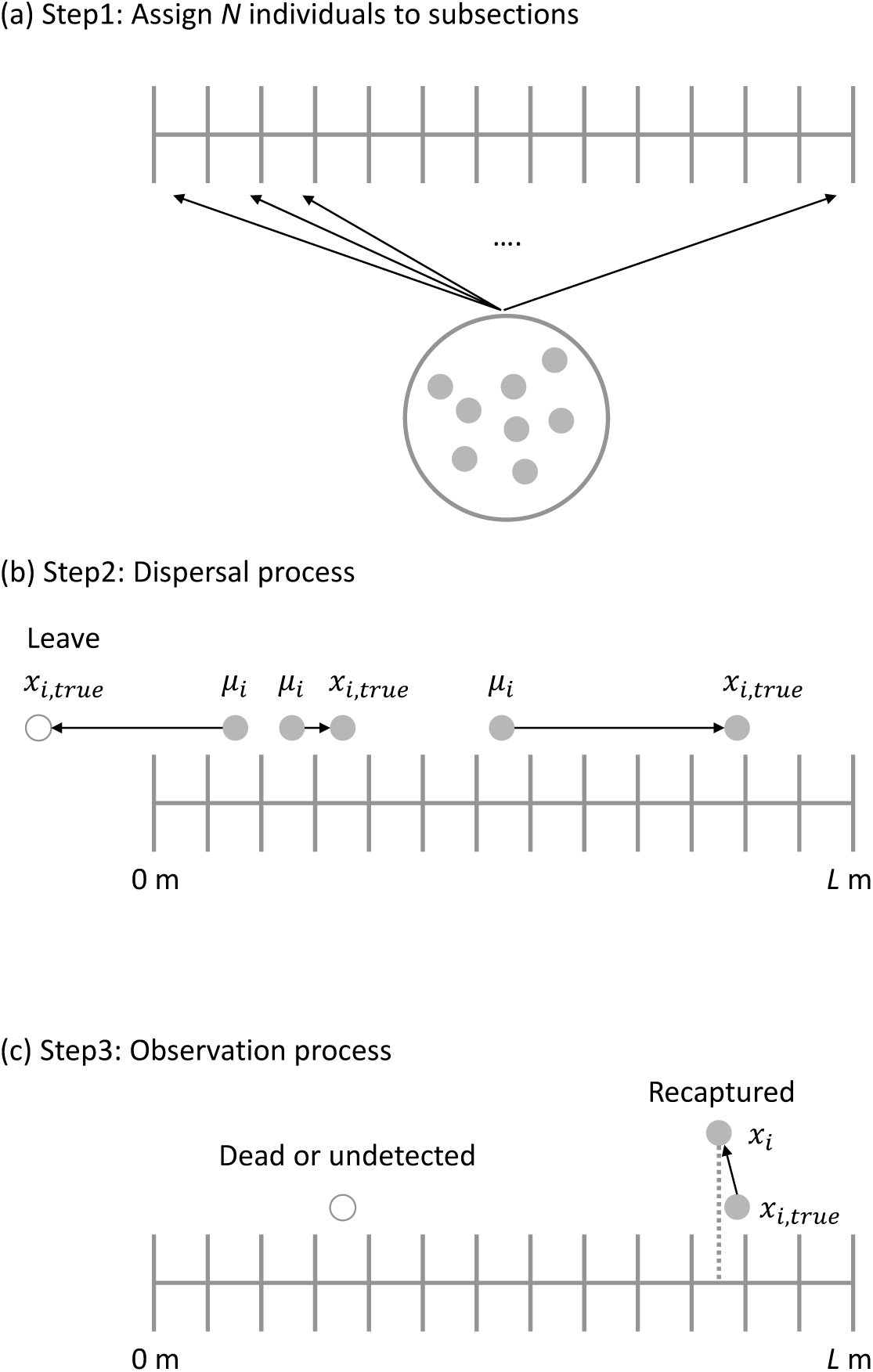
Procedure used to generate test datasets. (a) In step 1, *N* individuals were randomly distributed in the observation section, resembling a marking process in the field. (b) In step 2, marked individuals move freely from the center of the released subsection (*μ*_*i*_) according to a known dispersal kernel (in this case, a Laplace distribution with a mean dispersal distance *δ*). Since the observation section is a finite domain, individuals can leave the observation section and may never be recaptured (open circle in the figure). Individuals were considered to remain in the observation section when true location after dispersal *x*_*i,true*_was within a range of 0 – *L* m. (c) In step 3, individuals that remained in the observation section were subject to an incomplete recapture survey. Remained individuals were recaptured with recapture probability *ϕ*, and recapture location *x*_*i*_ was recorded as the center of the recapture subsection (the vertical dotted line in the figure). A certain proportion of remained individuals (1 − *ϕ*) may not be recaptured because they may have died or undetected (open circle in the figure).

I estimated average dispersal distance *δ* and recapture probability *ϕ* using the DOCM and a simple dispersal model. The simple dispersal model is a “control” that does not model the observation process and the average dispersal distance was estimated as *x*_*i*_|*μ*_*i*_, *δ ~ Laplace*(*μ*_*i*_, *δ*). The estimates of average dispersal distances were compared between the models. Meanwhile, the estimated recapture probability *ϕ* was compared with the proportion of individuals recaptured (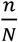, where *n* is the number of recaptured individuals) in the test dataset used to estimate *ϕ* because the simple dispersal model does not estimate *ϕ*. Finally, I assessed the accuracy (i.e., the closeness of the median estimate to the true parameter) and precision (i.e., the 95% credible interval [CI]) of the estimated parameters.

The Bayesian models were fitted to the test datasets to estimate *δ* and *ϕ*. Vague priors were assigned to the parameters: a half-Cauchy distribution for δ (scale = 500) and a uniform distribution for *ϕ* (range: 0 – 1). Three Markov chain Monte Carlo (MCMC) chains were run with 4500 iterations, 1500 burn-ins, and 3 thin numbers, resulting in a total of 1500 MCMC samples. Convergence was assessed by whether the R–hat indicator of each parameter had reached a value near 1. All statistical analysis was conducted using R 3.5.1 (RCoreTeam 2019) and JAGS 4.3.0 (Plummer 2003). A sample of JAGS scripts for the DOCM was provided in Box 1. R and JAGS scripts used in simulations will be made available at Github upon publication.

#### Box 1

Sample JAGS script for the dispersal-observation coupled model

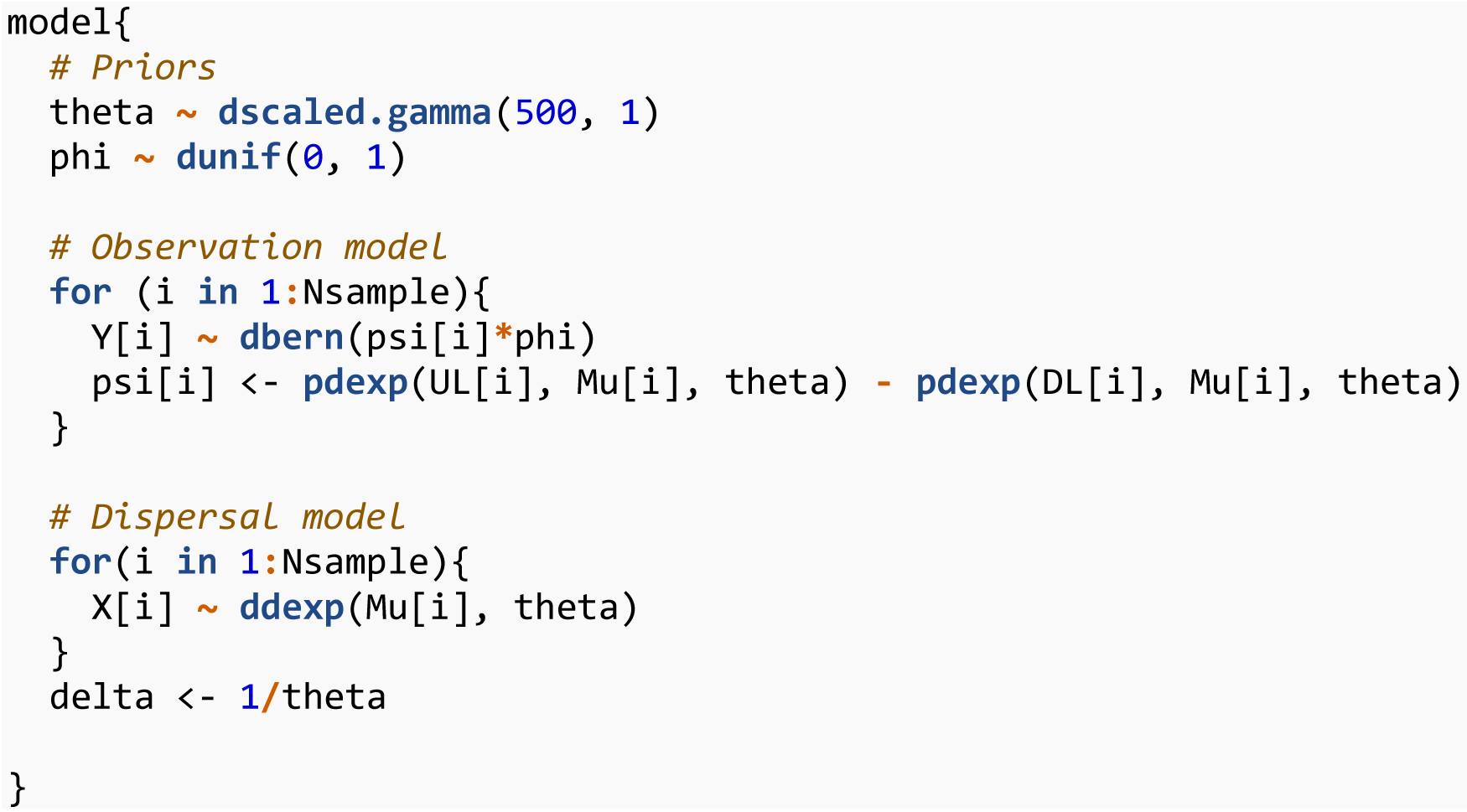

## Results and discussion

### Model performance

The DOCM performed well under various sampling designs. Figure 4 shows the relationship between the true and estimated values of δ (denoted as *δ*_*true*_ and *δ*_*est*_, respectively) when the recapture probability *ϕ* was 0.50. The parameter estimates from the DOCM were always closer to the true values (compare red and black lines in Figure 4) compared with those derived from the simple dispersal model without the observation process. The degree of improvement was significant. While 95% CIs of the simple dispersal model tended not to include *δ*_true_, the DOCM was more likely to encompass the true values especially when the observation length was long enough (*L* = 1000 m). Similarly, the DOCM provided less biased estimates of recapture probability *ϕ*, a composite of survival and detection probabilities (Figure 5). The estimated *ϕ* was higher than the proportion of individuals recaptured in the test dataset (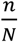, where *n* is the number of recaptured individuals) because it was corrected for permanent emigration. However, as the *δ*_*true*_ increases, the DOCM became underestimating the parameters, though the degree of bias is better than the simple dispersal model. This pattern was apparent when the observation section was short relative to *δ*_*true*_ and is caused by the substantial number of individuals leaving behind the observation section. For each of the parameters, these results were qualitatively similar irrespective of *ϕ*_*true*_, although higher values of *ϕ*_*true*_ led to the narrower range of 95% CIs for *δ*_*est*_ as more individuals were recaptured. Detailed results with different values of *ϕ*_*true*_ were provided in Figures S1 – S4.

**Figure 4.**
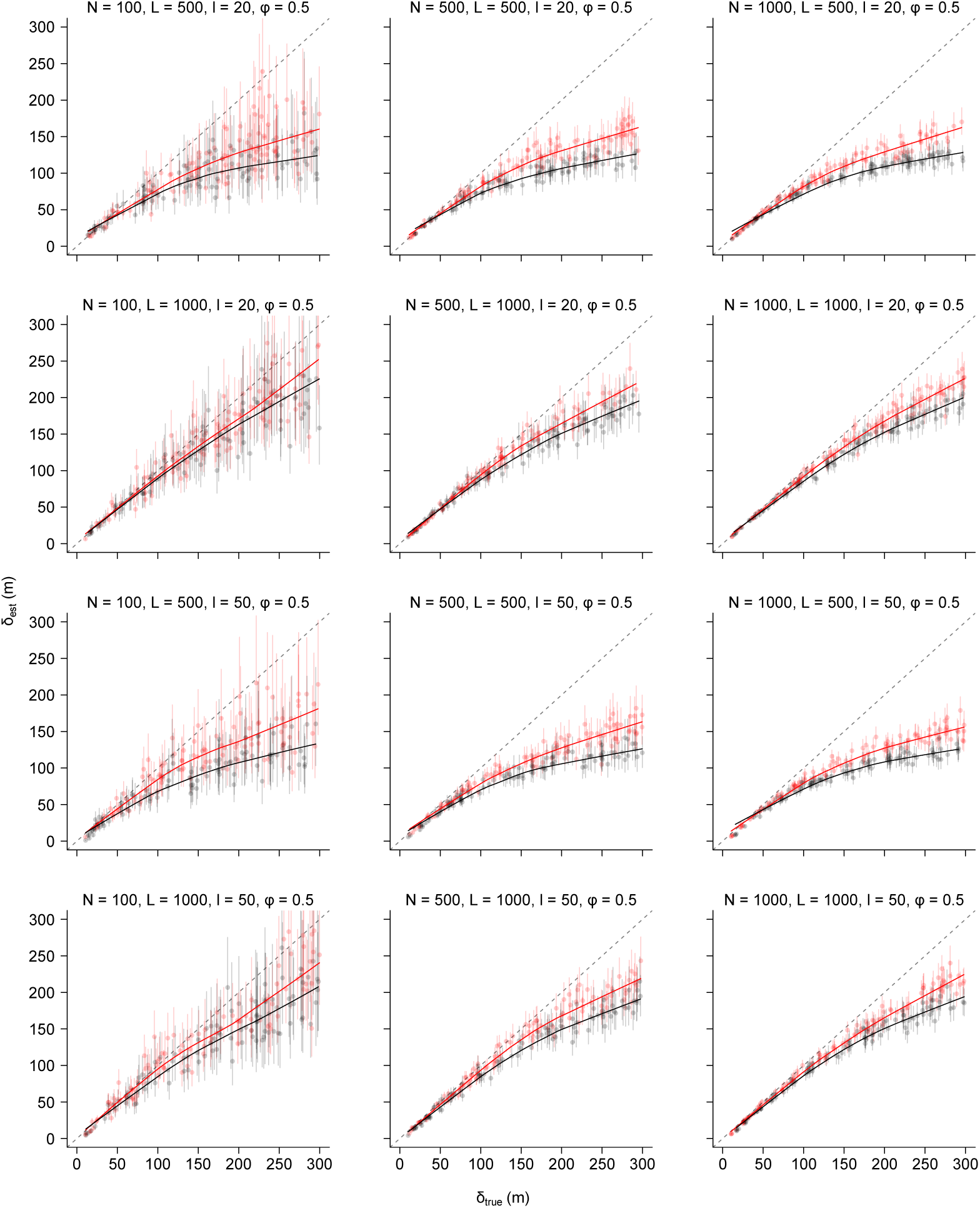
The relationship between true (*y*-axis) and estimated (*x*-axis) average dispersal distances when the true recapture probability *ϕ*_*true*_= 0.5. Red points are the median estimates from the DOCM, while grey points showing the median estimates from the simple dispersal model. Error bars are 95% credible intervals. Gray broken lines denote a 1:1 relationship. Different panels are estimates under different sampling designs, and the values of sampling design factors are shown on the top of each panel. Red and gray solid lines are smooths for the DOCM and the simple dispersal model, respectively.

**Figure 5.**
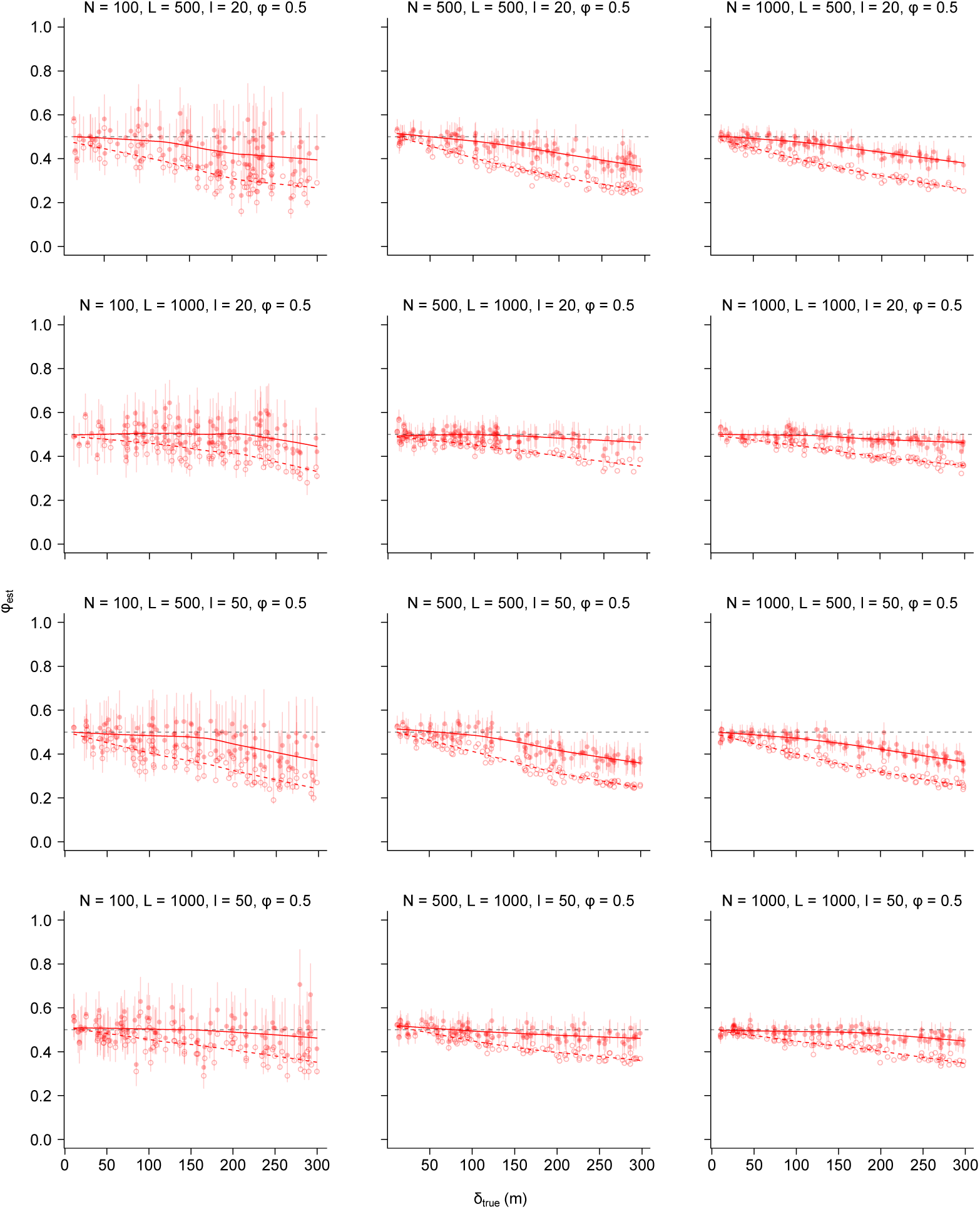
The relationship between the estimated recapture probability *ϕ*_*est*_ and true average dispersal distance *δ*_*true*_when the true recapture probability *ϕ*_*true*_= 0.5. Red filled points are the median estimates from the DOCM while red open points denoting the proportion of individuals recaptured for each test dataset. Error bars are 95% credible intervals. Grey broken lines denote the true recapture probability *ϕ*_*true*_. Different panels are estimates under different sampling designs, and the values of sampling design factors are shown on the top of each panel. Solid and broken red lines are smooths for the DOCM estimates and the proportion of individuals recaptured, respectively.

The number of individuals marked (*N*), observation section length (*L*) and spatial resolution (*l*) had distinct effects on *δ*_*est*_ and *ϕ*_*est*_. The number of individuals marked had a clear influence on the precision of the parameter estimates. The 95% CIs of *δ*_*est*_ and *ϕ*_*est*_ (error bars in Figures 4 and 5) became narrower clearly when *N* increased from 100 to 500 individuals. A further increase in *N*, however, did not improve the precision of the parameter estimates. Increasing *N* did not contribute to improving the accuracy of the parameter estimates (i.e., the closeness to *δ*_*true*_ and *ϕ*_*true*_). In contrast, the length of the observation section *L* was more influential on the accuracy of *δ*_*est*_ while having little influence on the precision of the parameter estimates (Figures 4 and 5). Increased *L* improved the accuracy of *δ*_*est*_ because long-distance dispersers were more likely to be recaptured. Neither accuracy nor precision was improved when the spatial resolution of sampling (smaller *l*) increased.

### Usefulness and limitations

The DOCM worked well under various sampling designs, proving its usefulness to infer dispersal processes in the wild. The DOCM can extract information from both recaptured and unrecaptured individuals, thereby improving the accuracy of parameter estimates. The DOCM, therefore, represents a promising tool to study dispersal processes. To apply the DOCM, users must obtain the following data: (1) individuals must be marked uniquely or by release subsection; (2) release location (*μ*_*i*_); (3) recapture location (*x*_*i*_); (3) spatial resolution of subsection length (*l*); (4) observation section length (*L*). These are a common dataset obtained through a mark-recapture study, so no additional work may be required to use the DOCM. Furthermore, if users have an independent estimate of detection probability *ξ* through multiple-pass removal (Dorazio *et al.* 2005) or other methods, it is also possible to estimate the true survival rate *s* that is corrected for permanent emigration (Terui *et al.* 2017). However, there are caveats when interpreting the results. As stated above, the estimated dispersal parameter *δ*_*est*_ and recapture probability *ϕ*_*est*_ can be biased when the average dispersal distance (*δ*_*est*_) exceeds ca. 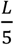. This happened because a significant portion of individuals may have left the observation section. Practically, the estimated dispersal parameter *δ*_*est*_ may be used to determine whether the average dispersal distance exceeds the threshold. In cases where 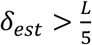, users shall acknowledge the potential bias in parameter estimates.

It is important to emphasize that different design factors (*L*, *N*, *l*) had different effects on the parameter estimates, corroborating the previous findings by Pépino *et al.* (2016). The results indicate that increasing the length of the observation section is most effective to increase the estimation accuracy of model parameters (the closeness to the true value). This is reasonable because increasing the length of the observation section is the only way to catch long-distance dispersers. In contrast, increasing the number of individuals marked is more important to improve the precision of the dispersal parameters (i.e., the range of 95% CI). Therefore, I recommend users paying close attention to the length of the observation section *L* and the number of individuals marked *N* when designing a mark-recapture study. Spatial resolution *l* had minimal influence on the accuracy and precision of parameter estimates, so this design component may be determined based on the biology of a study species.

### Model extension

The DOCM can be extended in two distinct ways. First, the dispersal model can capture the further complexity of the dispersal process. A growing body of evidence suggests that populations are composed of “resident” and “mobile” individuals with different behavioral and/or phenotypic characteristics (Clobert *et al.* 2012; Clobert *et al.* 2009; Cote *et al.* 2008; Cote *et al.* 2011; Cote *et al.* 2013; Terui *et al.* 2017). Such a linkage between dispersal and individual-level traits can be modeled using the following expression:

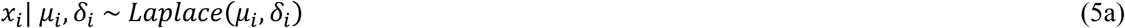

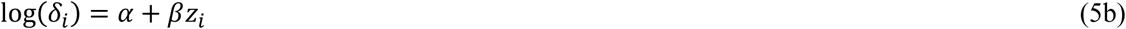

where *α* is the intercept, *β* the regression coefficient and *z*_*i*_ the linear predictor representing an individual trait. Expressed differently, equation 5 can be written as:

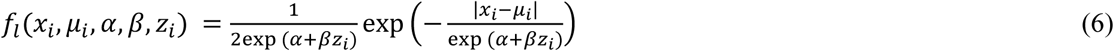

This model connects the trait variable *z*_*i*_ with the dispersal parameter *δ*_*i*_ by estimating *α* and *β*. In this model, individuals follow different dispersal kernels according to their ecological trait(s), such as body size. If the variable *z* is a random variable that follows a normal distribution with a mean *μ*_*z*_ and standard deviation *σ*_*z*_, *g*(*z*, *μ*_*z*_, *σ*_*z*_), then the composite dispersal kernel *h*(*x*_*i*_, *μ*_*i*_, *α*, *β*) is:

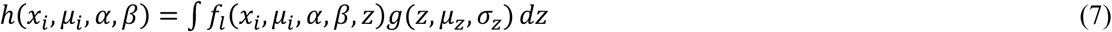

If *z* is a binary variable drawn from a Bernoulli distribution with a success probability *p*, the composite dispersal kernel is:

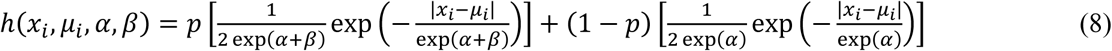

Equations 7 and 8 can be interpreted as an extension of a mixture Laplace dispersal kernel, in which a certain proportion of individuals are assigned randomly to a resident or mobile component in the model (Rodriguez 2002). The difference with a mixture Laplace dispersal kernel is that the above equations are explicit regarding “who is resident or mobile” as the expected dispersal distance for individual *i* (*δ*_*i*_) is related to ecological traits via the regression parameters. Terui *et al.* (2017) used this modeling framework to assess the effects of parasite infection on the dispersal of a stream fish species. It is important to note that there are many other dispersal kernels, such as a mixture of Gaussian dispersal kernels (Comte & Olden 2018; Nathan *et al.* 2012; Skalski & Gilliam 2000). Users may choose appropriate dispersal kernels given the ecology of a study species. I also point interested readers to Nathan *et al.* (2012) for dispersal kernels in two dimensional systems as another extension of the dispersal model.

Second, the observation model can also be extended to account for individual-level variability in recapture probability *ϕ*. Survival and detection probabilities may vary among individuals and ignoring this complexity could cause biased estimates of dispersal parameters. The simplest way to account for the variability is to model *ϕ*_*i*_ as a random variable drawn from a Beta distribution:

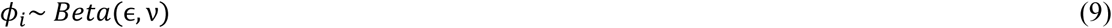

This allows the model to account for individual-level variation in recapture probability *ϕ*. If there are hypothesized predictors that could influence the recapture probability (e.g., habitat structure), such effects can be modeled as:

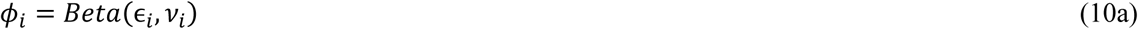

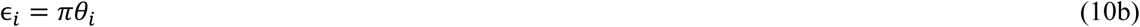

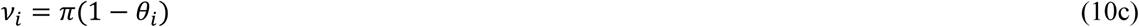

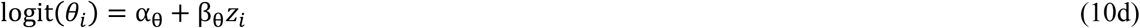

where *θ*_*i*_ is the expected recapture probability, *π* the dispersion parameter, *α*_*θ*_ the intercept and *β*_*θ*_ the regression coefficient. Therefore, the DOCM can deal with the complexity of field data.

### Conclusion

Dispersal is a fundamental process that drives the ecology and evolution of various organisms (Clobert *et al.* 2012) and quantifying dispersal is a critical task to forecast spatial dynamics of ecological systems (Hanski 1999; Hanski & Ovaskainen 2000; Terui *et al.* 2017). Although great strides have been made in how to quantify dispersal in the wild (e.g., genotyping) (Comte & Olden 2018; Morrissey & Ferguson 2011), direct measurements of dispersal still provide essential information for an understanding of spatial processes (Comte & Olden 2018; Kadoya & Inoue 2015; Terui *et al.* 2017). The Bayesian implementation of the DOCM provides extensive flexibility in the model formulation, offering a generic framework to study dispersal in the wild. Accurate inference of dispersal processes with sophisticated statistical models may enhance our ability to manage ecosystems in a changing world.

## Acknowledgments

I would like to express my sincere appreciation for all the help and kindness I have received from everyone who has supported my work.

## Supporting Information

**Figure S1.**
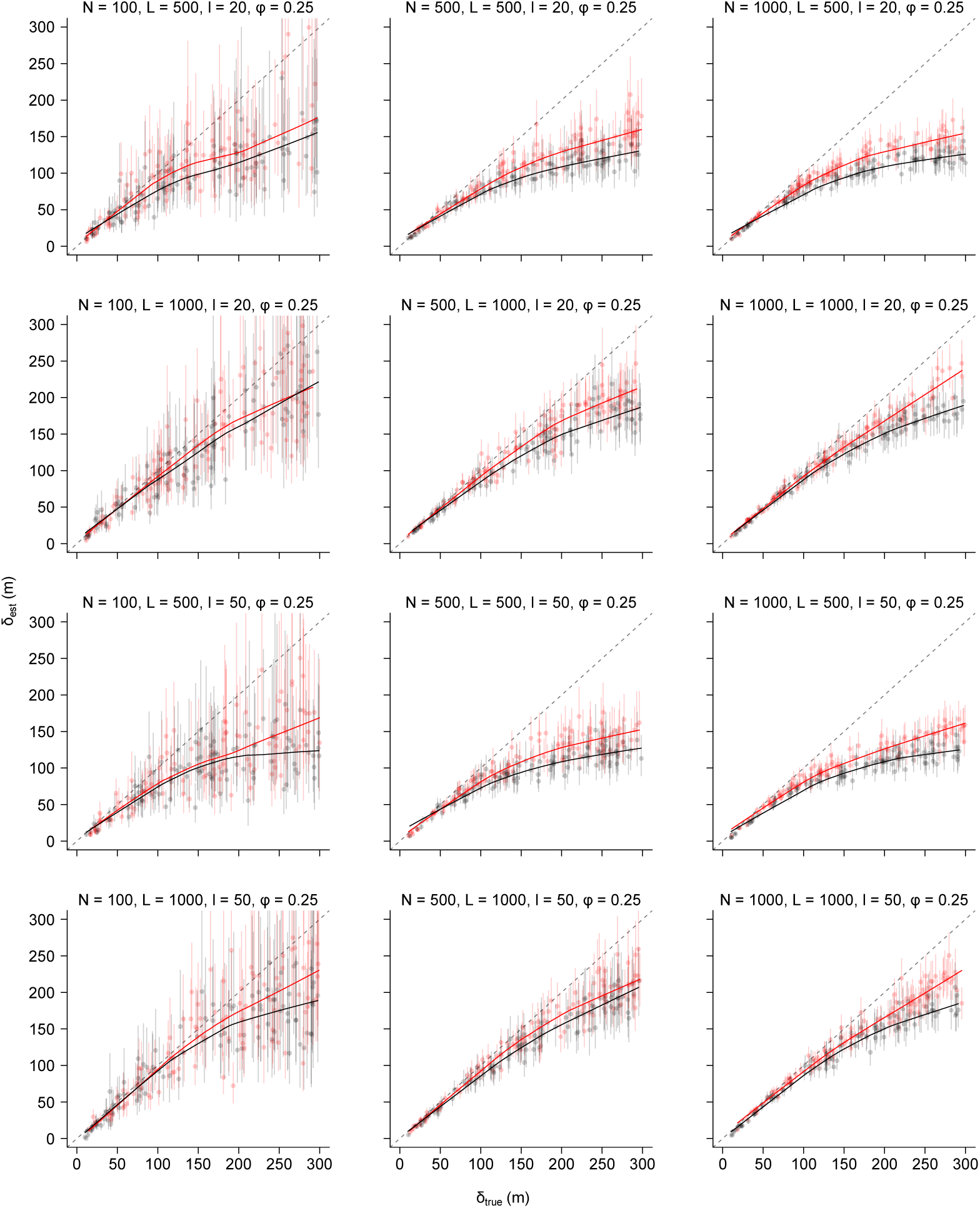
The relationship between true (*δ*_*true*_; *y*-axis) and estimated (*δ*_*est*_; *x*-axis) average dispersal distances when the true recapture probability *ϕ*_*true*_ = 0.25. Red points are the median estimates from the DOCM while grey points showing the median estimates from the simple dispersal model. Error bars are 95% credible intervals. Gray broken lines denote a 1:1 relationship. Different panels are estimates under different sampling designs, and the values of sampling design factors are shown on the top of each panel. Red and gray solid lines are smooths for the DOCM and the simple dispersal model, respectively.

**Figure S2.**
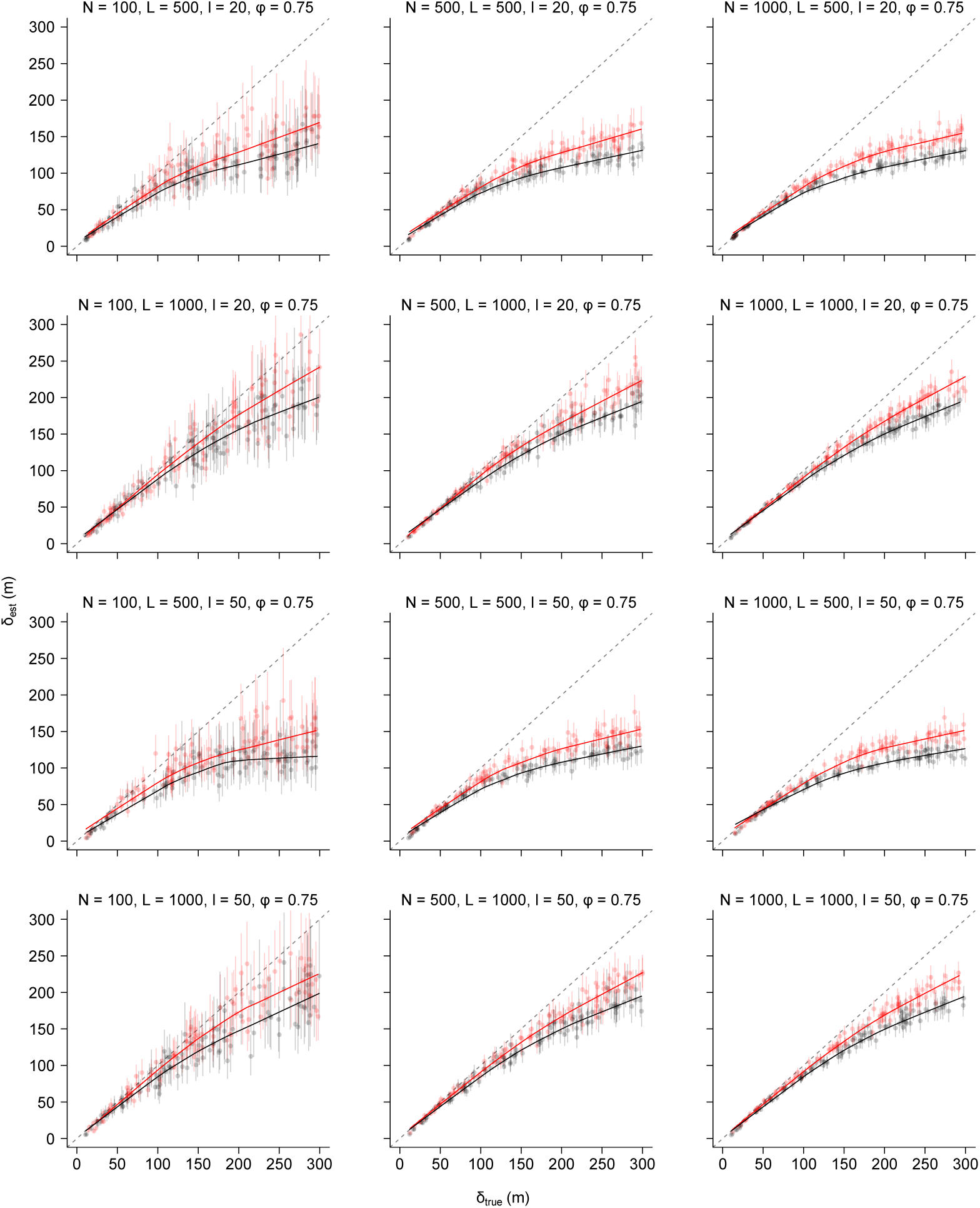
The relationship between true (*y*-axis) and estimated (*x*-axis) average dispersal distances when the true recapture probability *ϕ*_*true*_ = 0.75. See Figure S1 for captions.

**Figure S3.**
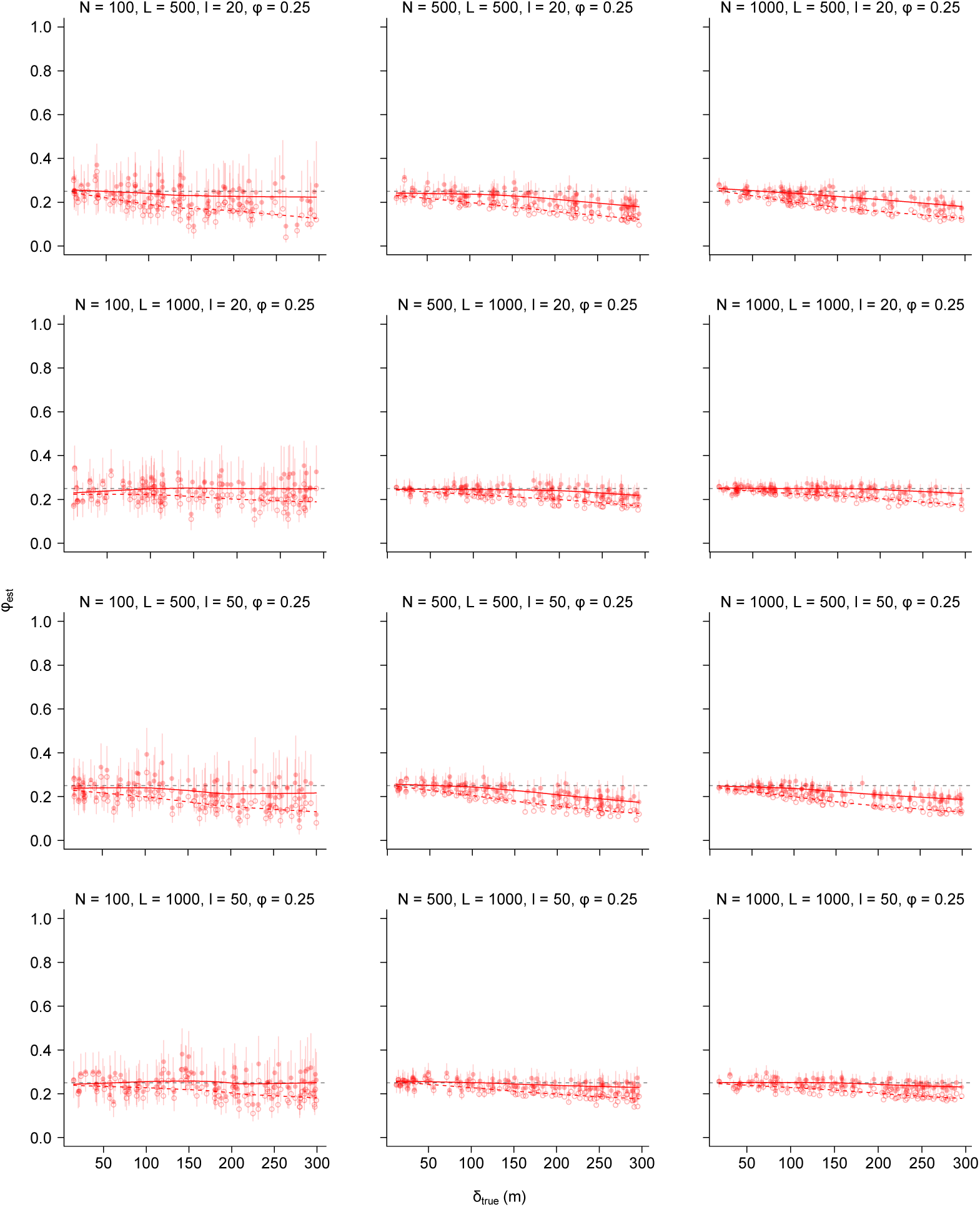
The relationship between the estimated recapture probability *ϕ*_*est*_ and true average dispersal distance *δ*_*true*_ when the true recapture probability *ϕ*_*true*_ = 0.25. Red filled points are the median estimates from the DOCM while red open points denoting the proportion of individuals recaptured for each test dataset. Error bars are 95% credible intervals. Grey broken lines denote the true recapture probability *ϕ*_*true*_. Different panels are estimates under different sampling designs, and the values of sampling design factors are shown on the top of each panel. Solid and broken red lines are smooths for the DOCM estimates and the proportion of individuals recaptured, respectively.

**Figure S4.**
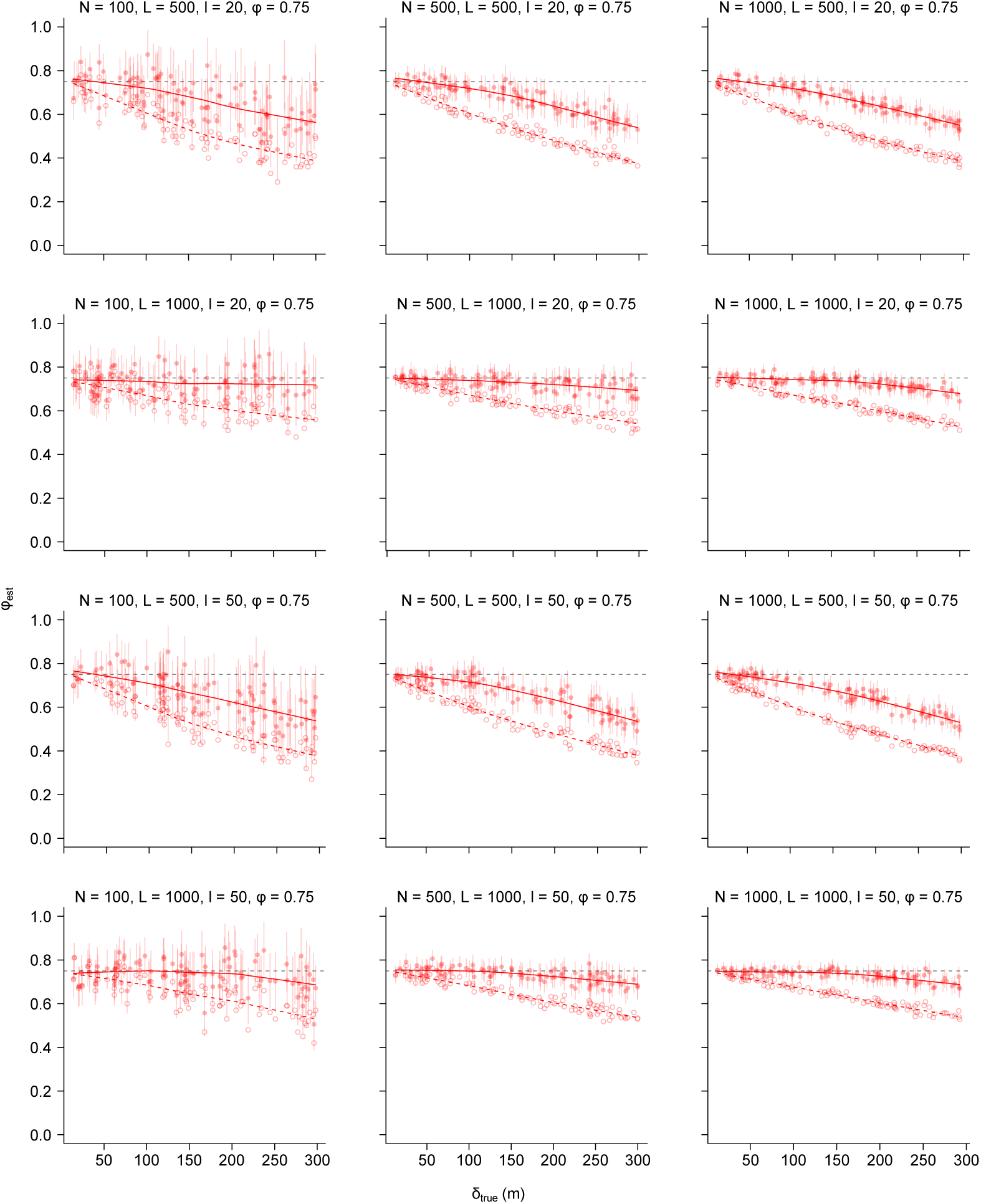
The relationship between the estimated recapture probability *ϕ*_*est*_ and true average dispersal distance *δ*_*true*_ when the true recapture probability *ϕ*_*true*_ = 0.75. See Figure S3 for captions.

